# A large-scale genome-wide enrichment analysis identifies new trait-associated genes, pathways and tissues across 31 human phenotypes^*^

**DOI:** 10.1101/160770

**Authors:** Xiang Zhu, Matthew Stephens

## Abstract

Genome-wide association studies (GWAS) aim to identify genetic factors that are associated with complex traits. Standard analyses test individual genetic variants, one at a time, for association with a trait. However, variant-level associations are hard to identify (because of small effects) and can be difficult to interpret biologically. “Enrichment analyses” help address both these problems by focusing on *sets of biologically-related variants*. Here we introduce a new model-based enrichment analysis method that requires only GWAS summary statistics, and has several advantages over existing methods. Applying this method to interrogate 3,913 biological pathways and 113 tissue-based gene sets in 31 human phenotypes identifies many previously-unreported enrichments. These include enrichments of the *endochondral ossification* pathway for adult height, the *NFAT-dependent transcription* pathway for rheumatoid arthritis, *brain-related* genes for coronary artery disease, and *liver-related* genes for late-onset Alzheimer’s disease. A key feature of our method is that inferred enrichments automatically help identify new trait-associated genes. For example, accounting for enrichment in *lipid transport* genes yields strong evidence for association between *MTTP* and low-density lipoprotein levels, whereas conventional analyses of the same data found no significant variants near this gene.

## Introduction

Genome-wide association studies (GWAS) have successfully identified many genetic variants – typically SNPs – underlying a wide range of complex traits [1–3]. GWAS are typically analyzed using “single-SNP” association tests, which assess the marginal correlation between the genotypes of each SNP and the trait of interest. This approach can work well for identifying common variants with sufficiently-large effects. However, for complex traits, most variants have small effects, making them difficult to identify even with large sample sizes [4]. Further, because many associated variants are non-coding it can be difficult to identify the biological mechanisms by which they may act.

Enrichment analysis – also referred to as “pathway” [5] or “gene set” [6] analysis – can help tackle both these problems. Instead of analyzing one variant at a time, enrichment analysis assesses groups of related variants. The idea – borrowed from enrichment analysis of gene expression [7] – is to identify groups of biologically-related variants that are “enriched” for associations with the trait: that is, they contain a higher fraction of associated variants than would be expected by chance. By pooling information across many genetic variants this approach has the potential to detect enrichments even when individual genetic variants fail to reach a stringent significance threshold [5]. And because the sets of variants to be analyzed are often defined based on existing biological knowledge, an observed enrichment automatically suggests potentially relevant biological processes or mechanisms.

Although the idea of enrichment analysis is simple, there are many ways to implement it in practice, each with its own advantages and disadvantages. Here we build on a previous approach [8] that has several attractive features not shared by most methods. These features include: it accounts for linkage disequilibrium (LD) among associated SNPs; it assesses SNP sets for enrichment directly, without requiring intermediate steps like imposing a significance cut-off or assigning SNP-level associations to specific genes; and it can re-assess (“prioritize”) variant-level associations in light of inferred enrichments to identify which genetic factors are driving the enrichment.

Despite these advantages, this approach has a major limitation: it requires individual-level genotypes and phenotypes, which are often difficult or impossible to obtain, especially for large GWAS meta analyses combining many studies. Our major contribution here is to overcome this limitation, and provide an implementation that requires only GWAS summary statistics (plus LD estimates from a suitable reference panel). This allows the method to be applied on a scale that would be otherwise impractical. Here we exploit this to perform enrichment analyses of 3,913 biological pathways and 113 tissue-based gene sets for 31 human phenotypes. Our results identify many novel pathways and tissues relevant to these phenotypes, as well as some that have been previously identified. By prioritizing variants within the enriched pathways we identify several trait-associated genes that do not reach genome-wide significance in conventional analyses of the same data. The results highlighted here demonstrate the potential for these enrichment analyses to yield novel insights from existing GWAS summary data.

## RESULTS

**Method overview**. Figure 1 provides a schematic overview of the method. In brief, we combine the enrichment model from [8], with the multiple regression model for single-SNP association summary statistics from [9], to create a model-based enrichment method for GWAS summary data. We refer to the method as **R**egression with **S**ummary **S**tatistics, or RSS.

**Fig 1:**
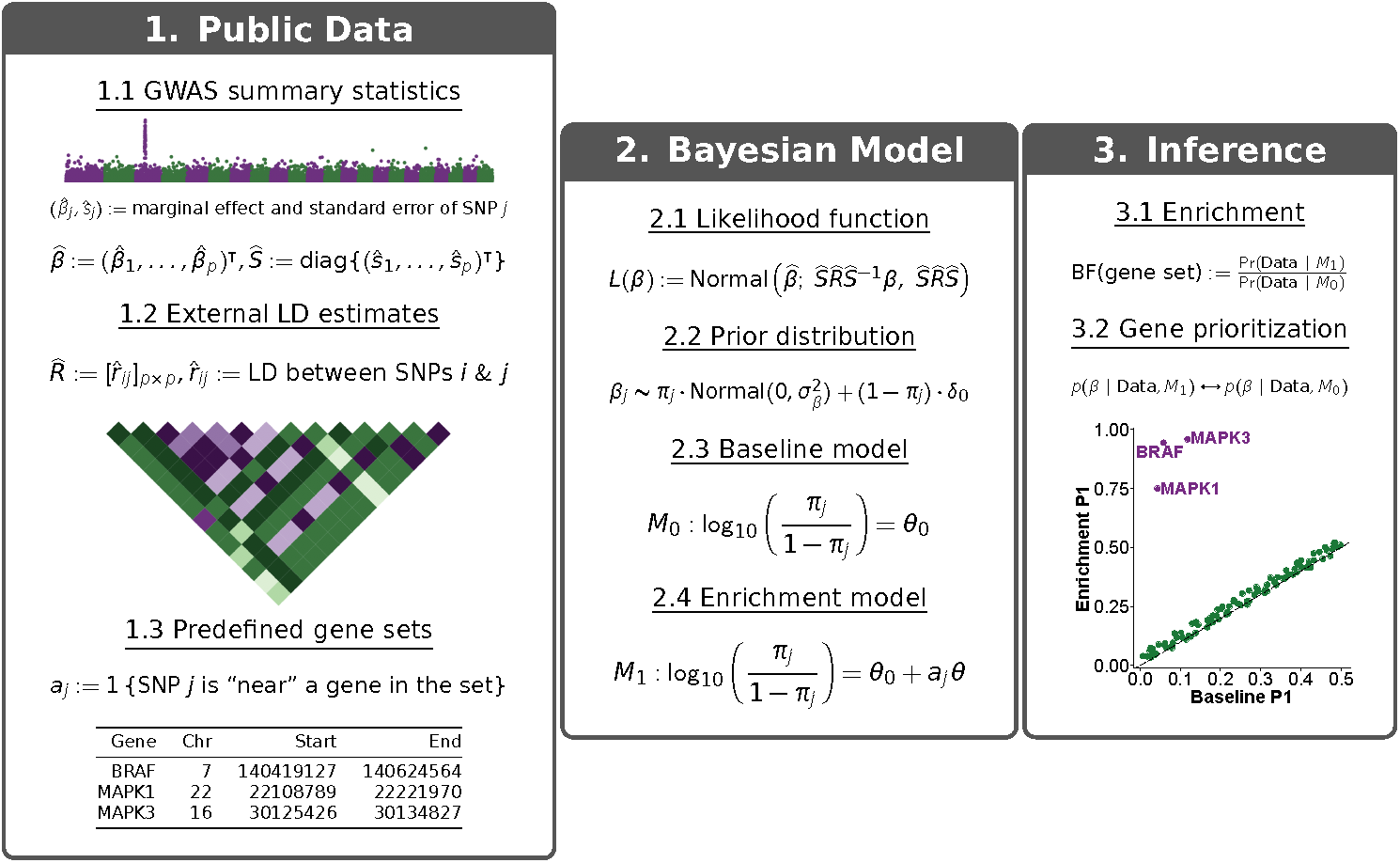
Schematic overview of RSS, a model-based enrichment analysis method for GWAS summary statistics. RSS combines three types of public data: GWAS summary statistics (1.1), external LD estimates (1.2), and predefined SNP sets (1.3). GWAS summary statistics consist of a univariate effect size estimate 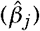 and corresponding standard error (*ŝ_j_*) for each SNP, which are routinely generated in GWAS. External LD estimates are obtained from an external reference panel with ancestry matching the population of GWAS cohorts. SNP sets here are derive from gene sets based on biological pathways or sequencing data. We combine these three types of data by fitting a Bayesian multiple regression (2.1-2.2) under two models about the enrichment parameter (*θ*): the “baseline model” (2.3) that each SNP has equal chance of being associated with the trait (*M*_0_: *θ* = 0), and the “enrichment model” (2.4) that SNPs in the SNP set are more often associated with the trait (*M*_1_: *θ* > 0). To test enrichment, RSS computes a Bayes factor (BF) comparing these two models (3.1). RSS also automatically prioritizes SNPs within an enriched set by comparing the posterior distributions of genetic effects (*β*) under *M*_0_ and *M*_1_, facilitating the discovery of new trait-associated genes (3.2). See Methods and Supplementary Note for additional details.

Specifically RSS requires single-SNP effect estimates and their standard errors from GWAS, and LD estimates from an external reference panel with similar ancestry to the GWAS cohort. Then, for any given set of SNPs, RSS estimates an “enrichment parameter”, *θ*, which measures the extent to which SNPs in the set are more often associated with the phenotype. This enrichment parameter is on a log-10 scale, so *θ* = 2 means that the rate at which associations occur inside the set is ~ 100 times higher than the rate of associations outside the set, whereas *θ* = 0 means that these rates are the same. When estimating *θ* RSS uses a multiple regression model to account for LD among SNPs. For example, RSS will (correctly) treat data from several SNPs that are in perfect LD as effectively a single observation, and not multiple independent observations. RSS ultimately summarizes the evidence for enrichment by a Bayes factor (BF) comparing the “enrichment model” (*M*_1_: *θ* > 0) against the “baseline model” (*M*_0_: *θ* = 0). RSS also provides posterior distributions of genetic effects under *M*_0_ and *M*_1_, and uses them to prioritize variants within enriched sets.

Although enrichment analysis could be applied to any SNP set, here we focus on SNP sets derived from “gene sets” such as biological pathways. Specifically, for a given gene set, we define a corresponding SNP set as the set of SNPs within ±100 kb of the transcribed region of any member gene; we refer to such SNPs as “inside” the gene set. If a gene set plays an important role in a trait then genetic associations may tend to occur more often near these genes than expected by chance; our method is designed to detect this signal.

To facilitate large-scale analyses, we designed an efficient, parallel algorithm implementing RSS. Our algorithm exploits variational inference [10], banded matrix approximation [11] and an expectation-maximization accelerator [12]. Software is available at https://github.com/stephenslab/rss.

**Method comparison based on simulations**. The novelty of RSS lies in its use of whole-genome association summary statistics to infer enrichments, and more importantly, its automatic prioritization of genes in light of the inferred enrichments. We are not aware of any published method with similar features. However, there are methods that can learn either enrichments or gene-level associations from GWAS summary statistics, *but not both*. We compared RSS to them through simulations using real genotypes [13].

To benchmark its enrichment component, we compared RSS with a suite of conventional pathway methods, Pascal [15], and a polygenic approach, LDSC [14]. We started with simulations without model mis-specification, where “baseline” and “enrichment” datasets were generated from corresponding-models (*M*_0_ and *M*_1_). Figure 2a and Supplementary Figure 1 show the trade-off between false and true enrichment discoveries for each method. All methods are powerful when the true underlying genetic architecture is polygenic, whereas LDSC performs worse when the truth is sparse. In both polygenic and sparse scenarios RSS is the most powerful method.

**Fig 2:**
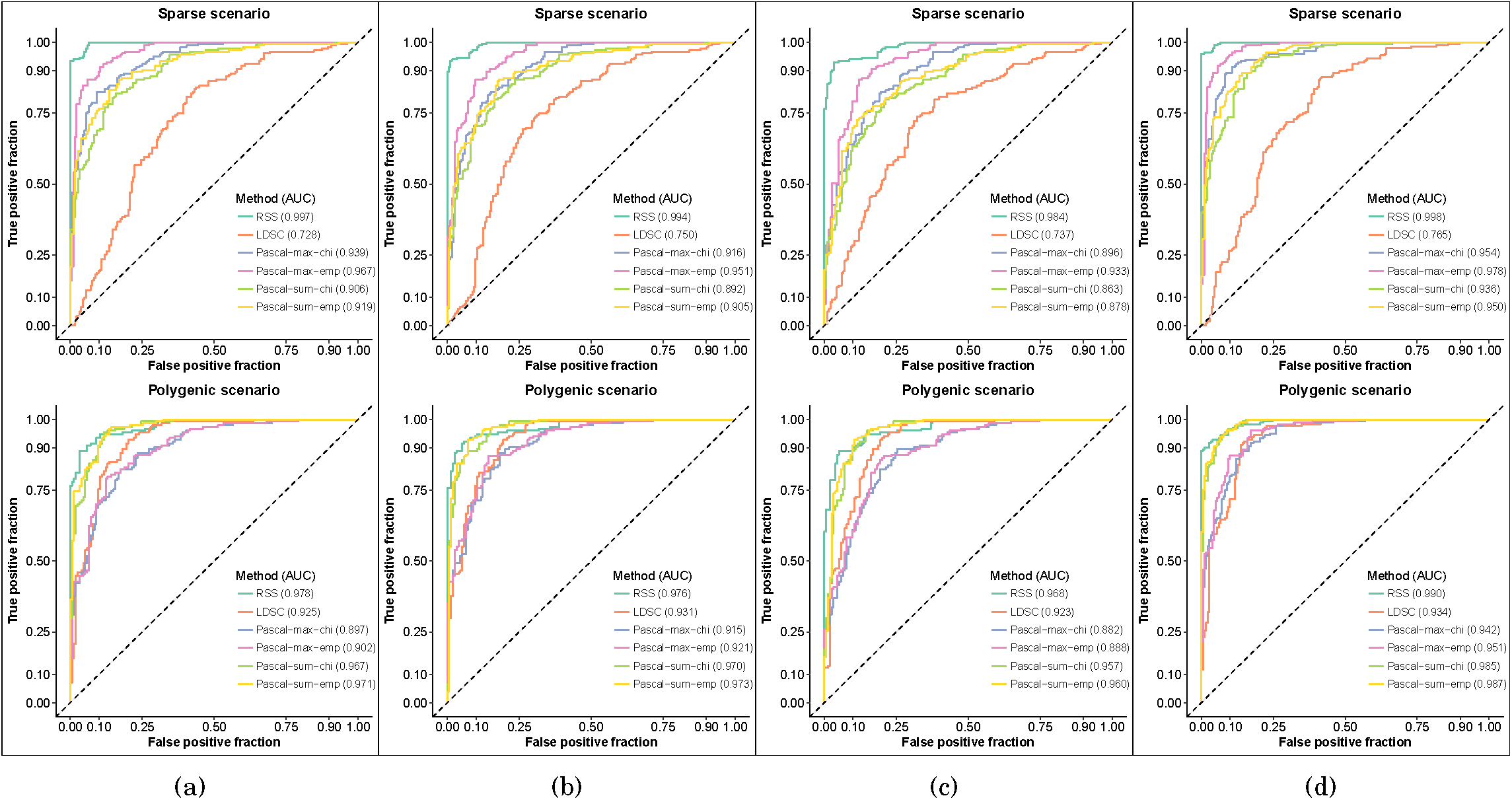
Comparison of RSS to other methods for identifying enrichments from GWAS summary statistics. We used real genotypes [13] to simulate individual-level data under two genetic architectures (“sparse” and “polygenic”) with four baseline-enrichment patterns: **(a)** baseline and enrichment datasets followed baseline (*M*_0_) and enrichment (*M*_1_) models in RSS; **(b)**baseline datasets assumed that a random set of near-gene SNPs were enriched for genetic associations and enrichment datasets followed *M*_1_; **(c)** baseline datasets assumed that a random set of coding SNPs were enriched for genetic associations and enrichment datasets followed *M*_1_; **(d)** baseline datasets followed *M*_0_ and enrichment datasets assumed that trait-associated SNPs were both more frequent, and had larger effects, inside than outside the target gene set. We computed the corresponding single-SNP summary statistics, and, on these summary data, we compared RSS with LDSC [14] and Pascal [15] using their default setups(URLs). Pascal includes two gene scoring options: maximum-of-*χ*^2^ (-max) and sum-of-*χ* ^2^(-sum), and two pathway scoring options:*χ* ^2^ approximation (-chi) and empirical sampling (-emp). For each simulated dataset, both Pascal and LDSC produced enrichment p-values, whereas RSS produced an enrichment BF; these statistics were used to rank the significance of enrichments. Each panel displays the trade-off between false and true enrichment discoveries for all methods in 200 baseline and 200 enrichment datasets of a given simulation scenario, and also reports the corresponding areas under the curve (AUCs), where a higher value indicates better performance. Simulation details and additional results are provided in Supplementary Figures 1-4.

Next, to assess its robustness to mis-specification, we performed three sets of simulations where either the baseline (*M*_0_) or enrichment (*M*_1_) model of RSS were mis-specified. Specifically, we considered scenarios where i) baseline data contained enrichments of random near-gene SNPs (Fig. 2b, Supplementary Fig. 2); ii) baseline data contained enrichments of random coding SNPs (Fig. 2c, Supplementary Fig. 3); and iii) enrichment data contained enrichments of effect sizes (Fig. 2d, Supplementary Fig. 4). The results show that RSS is highly robust to model mis-specifications, and still consistently outperforms Pascal and LDSC.

Recent analyses of GWAS summary statistics (e.g. [14]) often focus on the HapMap Project Phase 3 (HapMap3) SNPs [16], even though summary statistics of the 1000 Genomes Project SNPs [17] are available in many GWAS. Although not required, we used this “SNP subsetting” strategy in data analyses to reduce computation (Methods). To assess the impact of “SNP subsetting”, we simulated data using all 1000 Genome SNPs, applied the enrichment methods to summary statistics of HapMap3 SNPs only, and compared HapMap3-based results with results of analyzing all 1000 Genome SNPs. For all methods, analysis with and without “SNP subsetting” produced similar results (Supplementary Fig. 5). The robustness to “SNP subsetting” is perhaps unsurprising, since all methods utilize LD information (in different ways) to capture the effects of potentially excluded causal variants. As above, RSS has the highest power in this set of simulations.

Finally, to benchmark its prioritization component, we compared RSS with four gene-based association methods [18–21]. Figure 3 and Supplementary Figures 6-7 show the power of each method to identify gene-level associations. RSS substantially outperforms existing methods even in the absence of enrichments, especially in the polygenic scenario. This is because RSS exploits a multiple regression framework [9] to learn the genetic architecture from data of all genes and assesses their effects jointly, whereas other methods implicitly assume a fixed, sparse architecture and only use data of a single gene to estimate its effect. When datasets contain enrichments, RSS automatically leverages them, which existing methods ignore, to further improve the power.

**Fig 3:**
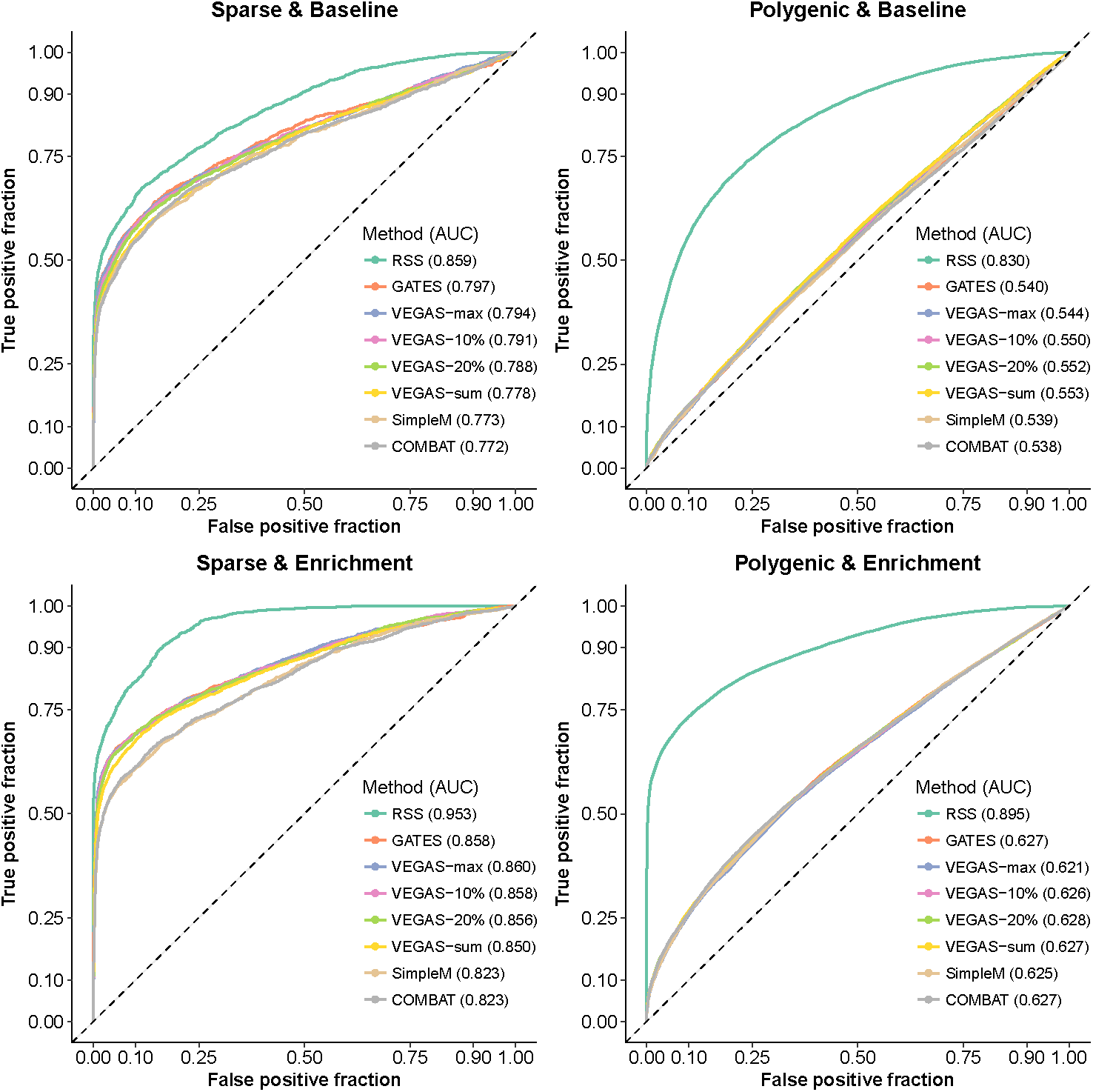
Comparison of RSS to other methods for identifying trait-associated genes from GWAS summary statistics. We used real genotypes [13] to simulate individual-level data under two genetic architectures (“Sparse” and “Polygenic”), with and without enrichment in the target gene set (“Enrichment” and “Baseline”), and then computed corresponding single-SNP summary statistics. On these summary data, we compared RSS with four other methods: SimpleM [18], VEGAS [19], GATES [20] and COMBAT [21]. We applied VEGAS to the full set of SNPs (-sum), to a specified percentage of the most significant SNPs (−10% and −20%), and to the single most significant SNP (-max), within ±100 kb of the transcribed region of each gene. All methods are available in the package COMBAT (URLs). For each simulated dataset, we defined a gene as “trait-associated” if at lease one SNP within ±100 kb of the transcribed region of this gene had non-zero effect. For each gene in each dataset, RSS produced the posterior probability that the gene was trait-associated. whereas the other methods produced association p-values; these statistics were used to rank the significance of gene-level associations. Each panel displays the trade-off between false and true gene-level associations for all methods in 100 datasets of a given simulation scenario, and reports the corresponding AUCs. Simulation details and additional results are provided in Supplementary Figures 6-7.

In conclusion, RSS outperforms existing methods in both enrichment and prioritization analysis, and is robust to a wide range of model mis-specification. To further investigate its real-world benefit, we applied RSS to analyze 31 complex traits and 4,026 gene sets.

**Multiple regression on 1.1 million variants across 31 traits**. The first step of our analysis is multiple regression of 1.1 million HapMap3 common SNPs for 31 traits, using GWAS summary statistics from 20,883-253,288 European ancestry individuals (Supplementary Table 1; Supplementary Fig. 8). This step essentially estimates, for each trait, a “baseline model” (*M*_0_) against which “enrichment models” (*M*_1_) can be compared. The fitted baseline model captures both the size and abundance (“polygenicity”) of the genetic effects on each trait, effectively providing a two-dimensional summary of the genetic architecture of each trait (Fig. 4a; Supplementary Fig. 9).

**Fig 4:**
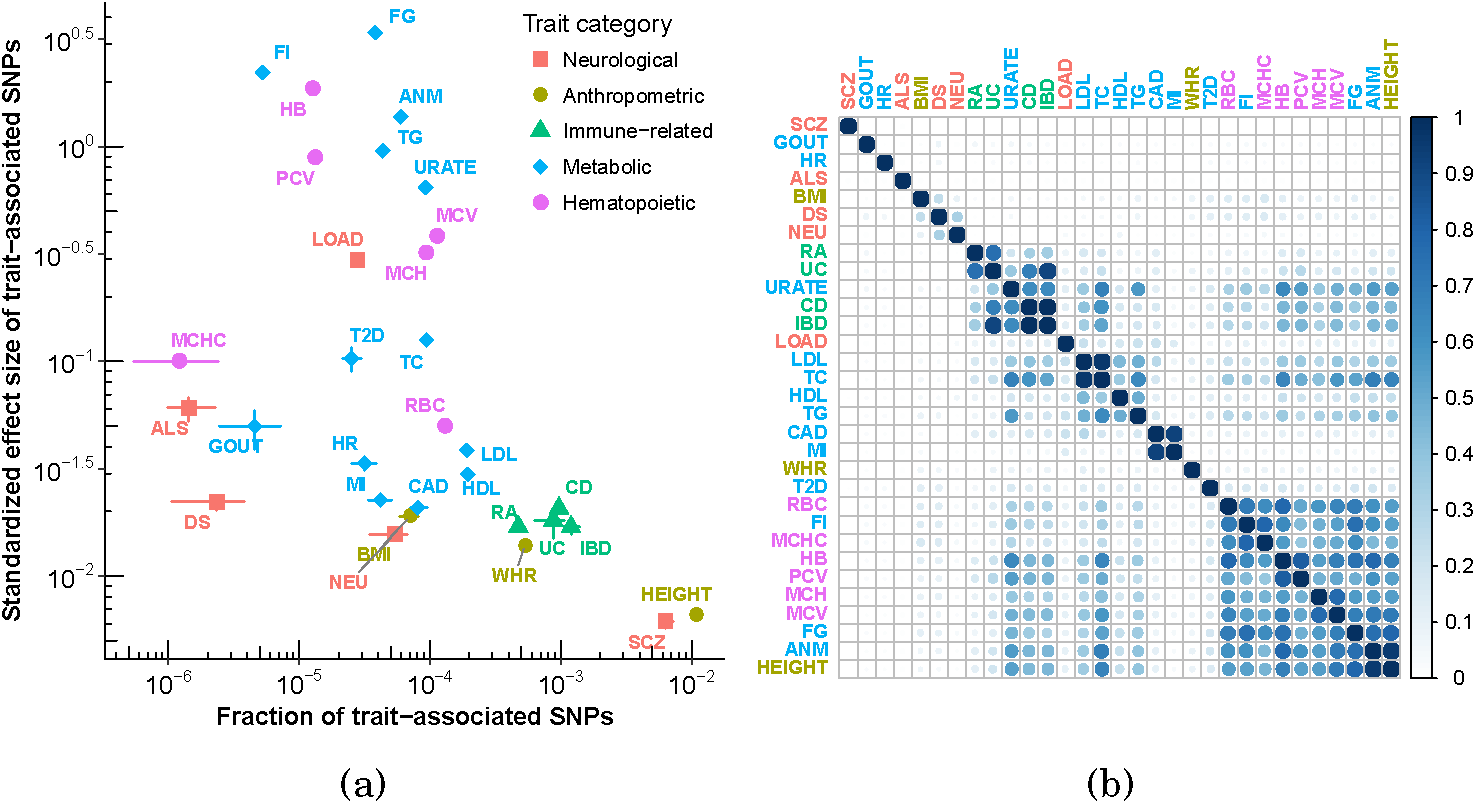
Baseline and enrichment analyses of 31 complex traits. For both panels, traits are colored by categories and labeled by abbreviations. **(a)** Summary of inferred effect size distributions of 31 traits based on HapMap3 SNPs. Results are from fitting the baseline model (*M*_0_) to 1.1 million common HapMap3 SNPs for each trait. We summarize effect size distribution using two numbers: the estimated fraction of trait-associated SNPs (average posterior probability of a HapMap3 SNP being associated with a trait; shown in *x*-axis) and the standardized effect size of trait-associated SNPs (average posterior mean effect size of all HapMap3 SNPs, normalized by the phenotypic standard deviation and the fraction of trait-associated SNPs; shown in *y*-axis). See Supplementary Note for details on these two quantities. Each dot represents a trait, with horizontal and vertical point ranges indicating posterior mean and 95% credible interval for each quantity. Note that some intervals are too small to be visible due to log-10 scales. See Supplementary Table 2 for numerical values of these intervals. **(b)** Pairwise sharing of 3,913 pathway enrichments among 31 traits. For each pair of traits, we estimated the proportion of pathways that are enriched in both traits, among pathways enriched in at least one of the traits (Methods). Traits are clustered by hierarchical clustering as implemented in the package corrplot (URLs). Darker color and larger shape represent higher sharing. The estimates of sharing are provided in Supplementary Table 3. ALS: amyotrophic lateral sclerosis [22]. DS: depressive symptoms [23]. LOAD: late-onset Alzheimer’s disease [24]. NEU: neuroticism [23]. SCZ: schizophrenia [25]. BMI: body mass index [26]. HEIGHT: adult height [27]. WHR: waist-to-hip ratio [28]. CD: Crohn’s disease [29]. IBD: inflammatory bowel disease [29]. RA: rheumatoid arthritis [30]. UC: ulcerative colitis [29]. ANM: age at natural menopause [31]. CAD: coronary artery disease [32]. FG: fasting glucose [33]. FI: fasting insulin [33]. GOUT: Gout [34]. HDL: high-density lipoprotein [35]. HR: heart rate [36]. LDL: low-density lipoprotein [35]. MI: myocardial infarction [32]. T2D: type 2 diabetes [37]. TC: total cholesterol [35]. TG: triglycerides [35]. URATE: serum urate [34]. HB: haemoglobin [38]. MCH: mean cell HB [38]. MCHC: MCH concentration [38]. MCV: mean cell volume [38]. PCV: packed cell volume [38]. RBC: red blood cell count [38].

The results emphasize that genetic architecture varies considerably among phenotypes: estimates of both polygenicity and effect sizes vary by several orders of magnitude. Height and schizophrenia stand out as being particularly polygenic, showing approximately 10 times as many estimated associated variants as any other phenotype. Along the other axis, fasting glucose, fasting insulin and haemoglobin show the highest estimates of effect sizes, with correspondingly lower estimates for the number of associated variants. Although not our main focus, these results highlight the potential for multiple regression models like ours to learn about effect size distributions and genetic architectures from GWAS summary statistics.

Fitting the baseline model also yields an estimate of effect size for each SNP. These can be used to identify trait-associated SNPs and loci. Reassuringly, these multiple-SNP results recapitulate many associations detected in single-SNP analyses of the same data (Supplementary Fig.s 10-12). For several traits, these results also identify additional associations (Supplementary Fig.s 13-14). These additional findings, while potentially interesting, may be difficult to validate and interpret. Enrichment analysis can help here: if the additional signals tend to be enriched in a plausible pathway, it may both increase confidence in the statistical results and provide some biological framework to interpret them.

**Enrichment analyses of 3,913 pathways across 31 traits**. We next performed enrichment analyses of SNP sets derived from 3,913 expert-curated pathways, ranging in size from 2 to 500 genes (Supplementary Fig.s 15-16). For each trait-pathway pair we computed a BF testing the enrichment model, and estimated the enrichment parameter *θ*.

Since these analyses involve large-scale computations that are subject to approximation error, we developed some “sanity checks” for confirming enrichments identified by RSS. Specifically these simple methods confirm that the z-scores for SNPs inside a putatively-enriched pathway have a different distribution from those outside the pathway (with more z-scores away from 0) – using both a visual check and a likelihood ratio statistic (Supplementary Fig. 17). Of note, these methods cannot replace RSS in the present study. The visual check requires human input, and thus is not suitable for large-scale analyses like ours. The likelihood ratio does not account for LD, and is expected to be less powerful (Supplementary Fig. 18).

**Table 1.** Top-ranked pathways for enrichment of genetic associations in complex traits. For each trait here we report the most enriched pathway (if any) that i) has an enrichment Bayes factor (BF) greater than 10^8^; ii) has at least 10 and at most 200 member genes; iii) has at least two member genes with *P*_1_ > 0.9 (denoted as “signals”) under the enrichment model; and iv) passes the visual and likelihood ratio sanity checks (Supplementary Fig. 17). All BFs reported here are larger than corresponding BFs that SNPs within ±100 kb of transcribed regions of all genes are enriched (Supplementary Fig. 19). The corresponding background and enrichment parameter estimates are provided in online results (URLs). *P*_1_: the posterior probability that at least one SNP within ±100 kb of the transcribed region of a given gene has non-zero effect on the target trait. CaN: calcineurin. NFAT: nuclear factor of activated T cells. IL-2R*β*: interleukin-2 receptor beta chain. p75(NTR): p75 neurotrophin receptor. SIRT1: Sirtuin 1. GSL: glycosphingolipid. PC: Pathway Commons [39]. BS: NCBI BioSystems [40]. *α*: The full pathway name is “transport of glucose and other sugars, bile salts and organic acids, metal ions and amine compounds”.

| Phenotype | Top enriched pathway | Database<br>(Repository) | # of signals<br>(# of genes) | $\log_{10}$ BF |
| --- | --- | --- | --- | --- |
| **Neurological traits** |  |  |  |  |
| Depressive symptoms | Eicosapentaenoate biosynthesis | HumanCyc (PC) | 2 (12) | 36.9 |
| Alzheimer's disease | Golgi associated vesicle biogenesis | Reactome (PC) | 3 (49) | 83.7 |
| **Anthropometric traits** |  |  |  |  |
| Adult height | Endochondral ossification | WikiPathways (BS) | 57 (65) | 68.9 |
| **Immune-related traits** |  |  |  |  |
| Crohn's disease | Inflammatory bowel disease | KEGG (BS) | 24 (61) | 25.6 |
| Inflammatory bowel disease | Inflammatory bowel disease | KEGG (BS) | 26 (61) | 24.2 |
| Rheumatoid arthritis | CaN-regulated NFAT-dependent transcription in lymphocytes | PID (BS) | 11 (45) | 10.0 |
| Ulcerative colitis | Inflammatory bowel disease | KEGG (BS) | 16 (61) | 11.8 |
| **Metabolic traits** |  |  |  |  |
| Age at natural menopause | IL-2Rβ in T cell activation | BioCarta | 2 (37) | 866.7 |
| Coronary artery disease | p75(NTR)-mediated signaling | PID (BS) | 4 (55) | 16.0 |
| Fasting glucose | Hexose transport | Reactome (BS) | 4 (47) | 1,898.4 |
| Gout | Osteoblast signaling | WikiPathways (BS) | 2 (13) | 30.6 |
| High-density lipoprotein | Statin pathway | WikiPathways (BS) | 18 (30) | 113.9 |
| Low-density lipoprotein | Chylomicron-mediated lipid transport | Reactome (PC) | 11 (17) | 65.5 |
| Myocardial infarction | Glutathione synthesis and recycling | Reactome (PC) | 2 (11) | 9.6 |
| Total cholesterol | Glucose transport | Reactome (BS) | 2 (41) | 833.2 |
| Triglycerides | Validated targets of C-MYC transcriptional activation | PID (BS) | 3 (79) | 604.9 |
| Serum urate | Transport of glucose and others<sup>a</sup> | Reactome (PC) | 4 (95) | 1,558.1 |
| **Hematopoietic traits** |  |  |  |  |
| Haemoglobin (HB) | RNA polymerase I transcription | Reactome (BS) | 27 (107) | 2,641.3 |
| Mean cell HB (MCH) | Meiotic synapsis | Reactome (PC) | 21 (72) | 2,334.3 |
| MCH concentration | SIRT1 negatively regulates ribosomal RNA expression | Reactome (PC) | 3 (63) | 700.8 |
| Mean cell volume | DNA methylation | Reactome (PC) | 28 (61) | 2,077.3 |
| Packed cell volume | RNA polymerase I promoter opening | Reactome (PC) | 27 (59) | 217.5 |
| Red blood cell count | GSL biosynthesis (neolacto series) | KEGG (PC) | 2 (21) | 391.2 |

Since genic regions may be generally enriched for associations compared with non-genic regions, we confirmed that top-ranked pathways often showed stronger evidence for enrichment than did the set containing all genes (Supplementary Fig. 19). We also created “null” (non-enriched) SNP sets by randomly drawing near-gene SNPs, and performed enrichment analyses of these “null” sets on real GWAS summary data. Enrichment signals of these simulated genic sets are substantially weaker than the actual top-ranked sets (Supplementary Fig. 20). Further, to check whether observed enrichments could be driven by other functional annotations (e.g. coding), we computed the correlation between enrichment BFs and proportions of gene-set SNPs falling into each of 52 functional categories [14]. Among 1,612 trait-category pairs, we did not observe any strong correlation (median 7.3×10^−3^; 95% interval [−0.08,0.21]; Supplementary Fig. 21). Together, these results suggest that observed enrichments are unlikely to be artifacts driven by model mis-specification.

For most traits our analyses identify many pathways with strong evidence for enrichment – for example, at a conservative threshold of BF ≥ 10^8^, 20 traits are enriched in more than 100 pathways per trait (Supplementary Fig. 22). Although the top enriched pathways for a given trait often substantially overlap (i.e. share many genes), several traits show enrichments with multiple non-overlapping or minimally-overlapping pathways (Supplementary Fig. 23). Table 1 gives examples of top enriched pathways, with full results available online (URLs).

Our results highlight many previously reported trait-pathway links. For example, the *Hedgehog pathway* is enriched for associations with adult height (BF=1.9 × 10^40^), consistent with both pathway function [41] and previous GWAS [27]. Other examples include *interleukin-23 mediated signaling* with inflammatory bowel disease (BF=3.1×10^23^; [42]), *T helper cell surface molecules* with rheumatoid arthritis (BF=3.2 × 10^8^; [30]), *statin pathway* with levels of high-density lipoprotein cholesterol (BF=8.4 × 10^113^; [43]), and *glucose transporters* with serum urate (BF=1.2 × 10^1,558^; [34]).

The results also highlight several pathway enrichments that were not reported in corresponding GWAS publications. For example, the top pathway for rheumatoid arthritis is *calcineurin-regulated nuclear factor of activated T cells (NFAT)-dependent transcription in lymphocytes* (BF=1.1×10^10^). This result adds to the considerable existing evidence linking NFAT-regulated transcription to immune function [44] and bone pathology [45]. Other examples of novel pathway enrichments include *endochondral ossification* with adult height (BF=7.7 × 10^68^; [46]), *p75 neurotrophin receptor-mediated signaling* with coronary artery disease (BF=9.6 × 10^15^; [47]), and *osteoblast signaling* with gout (BF=3.8 × 10^30^; [48]).

**Overlapping pathway enrichments among related traits**. Some pathways show enrichment in multiple traits. To gain a global picture of shared pathway enrichments among traits we estimated the proportions of shared pathway enrichments for all pairs of traits (Fig. 4b; Methods). Clustering these pairwise sharing results highlights four main clusters of traits: immune-related diseases, blood lipids, heart disorders and red blood cell phenotypes. Blood cholesterol shows strong pairwise sharing with serum urate (0.67), haemoglobin (0.66) and fasting glucose (0.53), which could be interpreted as a set of blood elements. Serum urate shows moderate to strong sharing with rheumatoid arthritis (0.19) and inflammatory bowel diseases (0.38-0.63), possibly due to the function of urate crystals in immune responses [49] Further, Alzheimer’s disease shows moderate sharing with blood lipids (0.17-0.23), heart diseases (0.15-0.21) and inflammatory bowel diseases (0.10-0.13). This seems consistent with recent data linking Alzheimer’s disease to lipid metabolism [50], vascular disorder [51] and immune activation [52]. The biologically relevant clustering of shared pathway enrichments helps demonstrate the potential of large-scale GWAS data to highlight similarities among traits, complementing other approaches such as clustering of shared genetic effects [53] and co-heritability analyses [54].

**Novel trait-associated genes informed by enriched pathways**. A key feature of RSS is that pathway enrichments, once identified, are automatically used to “prioritize” associations at variants near genes in the pathway. Specifically, RSS gives almost identical estimates of the background parameter (*θ*_0_) in both baseline and enrichment analyses (Supplementary Fig. 24), and yields a positive estimate of the enrichment parameter (*θ*) if the pathway is identified as enriched (Supplementary Fig. 25). The positive estimate of *θ* increases the prior probability of association for SNPs in the pathway, which in turn increases the posterior probability of association for these SNPs.

This ability to prioritize associations, which is not shared by most enrichment methods, has several important benefits. Most obviously, prioritization analyses can detect additional genetic associations that may otherwise be missed. Furthermore, prioritization facilitates the identification of genes influencing a phenotype in two ways. First, it helps identify genes that may explain individual variant associations, which is itself an important and challenging problem [55]. Second, prioritization helps identify genes that drive observed pathway enrichments. This can be useful to check whether a pathway enrichment may actually reflect signal from just a few key genes, and to understand enrichments of pathways with generic functions.

To illustrate, we performed prioritization analyses on the trait-pathway pairs showing strongest evidence for enrichment. Following previous Bayesian GWAS analyses [8, 56], here we evaluated genetic associations at the level of loci, rather than individual SNPs. Specifically, for each locus we compute *P*_1_, the posterior probability that at least one SNP in the locus is associated with the trait, under both the baseline and enrichment hypothesis. Differences in these two *P*_1_ estimates reflect the influence of enrichment on the locus.

The results show that prioritization analysis typically increases the inferred number of genetic associations (Supplementary Fig. 26), and uncovers putative associations that were not previously reported in GWAS. For example, enrichment in *chylomicron-mediated lipid transport* pathway (BF=3.4 × 10^65^; Fig. 5a) informs a strong association between gene *MTTP* (baseline *P*_1_: 0.14; enrichment *P*_1_: 0.99) and levels of low-density lipoprotein (LDL) cholesterol (Fig. 5b). This gene is a strong candidate for harboring associations with LDL: *MTTP* encodes microsomal triglyceride transfer protein, which has been shown to involve in lipoprotein assembly; mutations in *MTTP* cause abetalipoproteinemia (OMIM: 200100), a rare disease characterized by permanently low levels of apolipoprotein B and LDL cholesterol; and *MTTP* is a potential pharmacological target for lowering LDL cholesterol levels [57]. However, no genome-wide significant SNPs near *MTTP* were reported in single-SNP analyses of either the same data [35] (Fig. 5c), or more recent data with larger sample size [58] (Fig. 5d).

**Fig 5:**
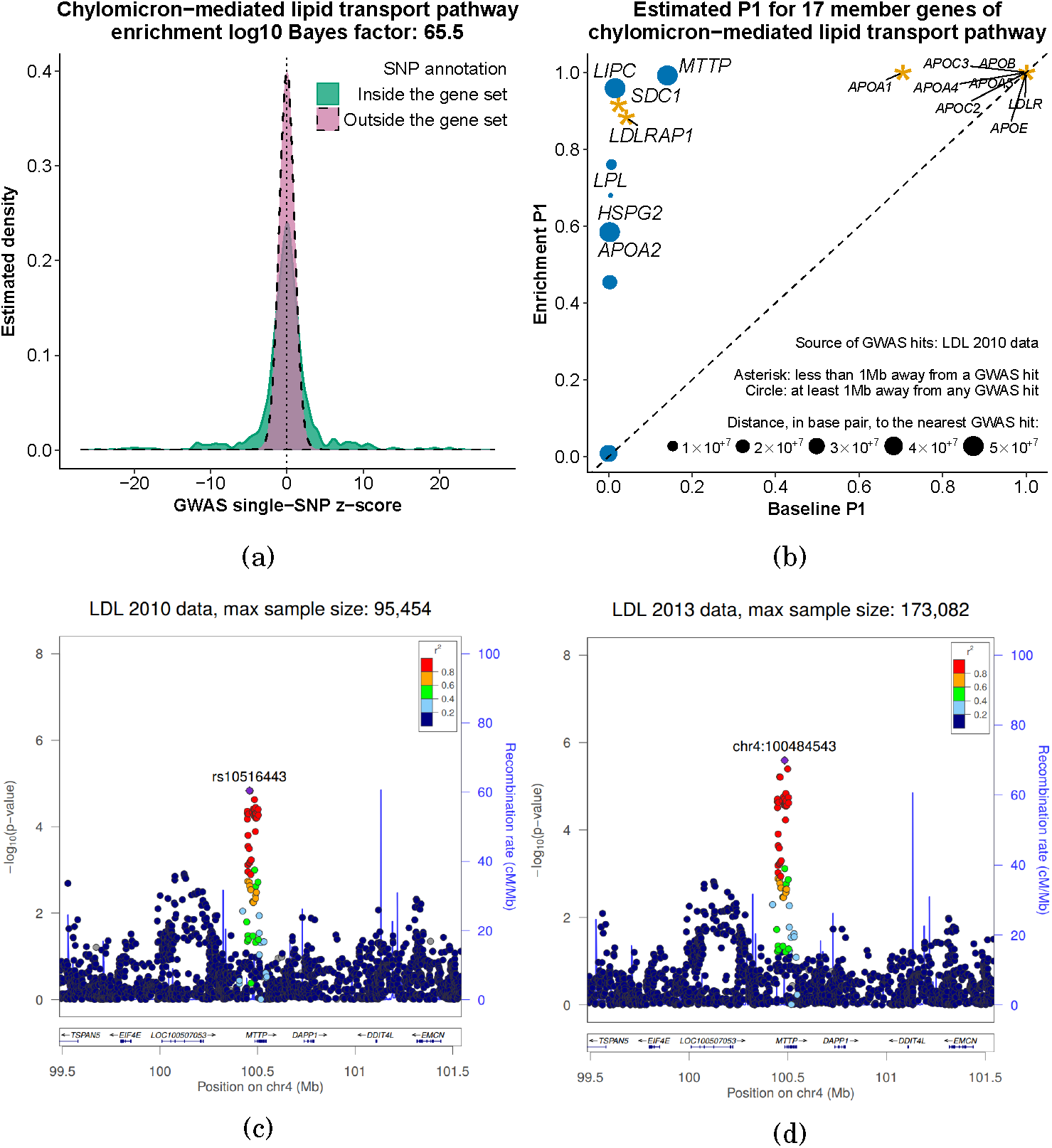
Enrichment of *chylomicron-mediated lipid transport* pathway informs a strong association between a member gene *MTTP* and levels of low-density lipoprotein (LDL) cholesterol. **(a)** Distribution of GWAS single-SNP z-scores from summary data published in 2010 [35], stratified by gene set annotations. The solid green curve is estimated from z-scores of SNPs within ± 100 kb of the transcribed region of genes in the *chylomicron-mediated lipid transport pathway* (“inside”), and the dashed reddish purple curve is estimated from z-scores of remaining SNPs (“outside”). This panel serves as a visual sanity check to confirm the observed enrichment. **(b)** Estimated posterior probability (*P*_1_) that there is at least one associated SNP within ± 100 kb of the transcribed region of each pathway-member gene under the enrichment model (*M*_1_) versus estimated *P*_1_ under the baseline model (*M*_0_). These gene-level *P*_1_ estimates and corresponding SNP-level statistics are provided in Supplementary Table 4. **(c)** Regional association plot for *MTTP* based on summary data published in 2010 [35]. **(d)** Regional association plot for *MTTP* based on summary data published in 2013 [58].

Prioritization analysis of the same *chylomicron-mediated lipid transport* pathway also yields several additional plausible associations (Fig. 5b). These include *LIPC* (baseline *P*_1_: 0.02; enrichment *P*_1_: 0.96) and *LPL* (baseline *P*_1_: 0.01; enrichment *P*_1_: 0.76). These genes play important roles in lipid metabolism and both reach genome-wide significance in single-SNP analyses of HDL cholesterol and triglycerides [35] although not for LDL cholesterol (Supplementary Fig. 27); and a multiple-trait, single-SNP analysis [59] of the same data also did not detect associations of these genes with LDL.

Several other examples of putatively novel associations that arise from our gene prioritization analyses, together with related literature, are summarized in Box 1.

### Box 1 Select putatively novel associations from prioritization analyses

**Adult height and *endochondral* ossification** (65 genes, log_10_BF = 68.9)

- *HDAC4* (baseline *P*_1_: 0.98; enrichment *P*_1_: 1.00) *HDAC4* encodes a critical regulator of chondrocyte hypertrophy during skeletogenesis [60] and osteoclast differentiation [61]. Haploinsufficiency of *HDAC4* results in chromosome 2q37 deletion syndrome (OMIM: 600430) with highly variable clinical manifestations including developmental delay and skeletal malformations.
- *PTH1R* (baseline *P*_1_: 0.94; enrichment *P*_1_: 1.00) *PTH1R* encodes a receptor that regulates skeletal development, bone turnover and mineral ion homeostasis [62]. Mutations in *PTH1R* cause several rare skeletal disorders (OMIM: 215045, 600002, 156400).
- *FGFR1* (baseline *P*_1_: 0.67; enrichment *P*_1_: 0.97) *FGFR1* encodes a receptor that regulates limb development, bone formation and phosphorus metabolism [63]. Mutations in *FGFR1* cause several skeletal disorders (OMIM: 101600, 123150, 190440, 166250).
- MMP13 (baseline *P*_1_: 0.45; enrichment *P*_1_: 0.93) *MMP13* encodes a protein that is required for osteocytic perilacunar remodeling and bone quality maintenance [64]. Mutations in *MMP13* cause a type of metaphyseal anadysplasia (OMIM: 602111) with reduced stature.

**IBD and *cytokine-cytokine receptor interaction*** (253 genes, log_10_BF = 21.3)

- *TNFRSF14* (a.k.a. *HVEM*; baseline *P*_1_: 0.98; enrichment *P*_1_: 1.00) *TNFRSF14* encodes a receptor that functions in signal transduction pathways activating inflammatory and inhibitory T-cell immune response. *TNFRSF14* expression plays a crucial role in preventing intestinal inflammation [65]. *TNFRSF14* is near a GWAS hit of celiac disease (rs3748816, *p* = 3.3×10^−9^) [66] and two hits of ulcerative colitis (rs734999, *p* = 3.3×10^−9^ [67]; rs10797432, *p* = 3.0×10^−12^ [68]).
- *FAS* (baseline *P*_1_: 0.82; enrichment *P*_1_: 0.99) *FAS* plays many important roles in the immune system [69]. Mutations in *FAS* cause autoimmune lymphoproliferative syndrome (OMIM: 601859).
- *IL6* (baseline *P*_1_: 0.27; enrichment *P*_1_: 0.87) *IL6* encodes a cytokine that functions in inflammation and the maturation of B cells, and has been suggested as a potential therapeutic target in IBD [70].

**CAD and p75(NTR)-*mediated signaling*** (55 genes, log_10_BF = 16.0)

- *FURIN* (baseline *P*_1_: 0.69; enrichment *P*_1_: 0.99) *FURIN* encodes the major processing enzyme of a cardiac-specific growth factor, which plays a critical role in heart development [71]. *FURIN* is near a *GWAS* hit (rs2521501 [72]) of both systolic blood pressure (*p* = 5.2×10^−19^) and hypertension (*p* = 1.9×10^−15^).
- *MMP3* (baseline *P*_1_: 0.43; enrichment *P*_1_: 0.97) A polymorphism in the promoter region of *MMP3* is associated with susceptibility to coronary heart disease-6 (OMIM: 614466). Inactivating *MMP3* in mice increases atherosclerotic plaque accumulation while reducing aneurysm [73].

**HDL and *lipid digestion, mobilization and transport*** (58 genes, log_10_BF = 89.8)

- *CUBN* (baseline *P*_1_: 0.24; enrichment *P*_1_: 1.00) *CUBN* encodes a receptor for intrinsic factor-vitamin B12 complexes (cubilin) that maintains blood levels of HDL [74]. Mutations in *CUBN* cause a form of congenital megaloblastic anemia due to vitamin B12 deficiency (OMIM: 261100). *CUBN* is near a GWAS hit of total cholesterol (rs10904908, *p* = 3.0×10^−11^ [58]).
- *ABCG1* (baseline *P*_1_: 0.01; enrichment *P*_1_: 0.89) *ABCG1* encodes an ATP-binding cassette transporter that plays a critical role in mediating efflux of cellular cholesterol to HDL [75].

**RA and *lymphocyte NFAT-dependent transcription*** (45 genes, log_10_BF = 10.0)

*PTGS2* (a.k.a. *COX2*; baseline *P*_1_: 0.74; enrichment *P*_1_: 0.98)
- *PTGS2*-specific inhibitors have shown efficacy in reducing joint inflammation in both mouse models [76] and clinical trials [77]. *PTGS2* is near a GWAS hit of Crohn’s disease (rs10798069, *p* = 4.3×10^−9^ [29])
- *PPARG* (baseline *P*_1_: 0.28; enrichment *P*_1_: 0.98)
- *PPARG* has important roles in regulating inflammatory and immune responses with potential applications in treating chronic inflammatory diseases including RA [78, 79].

### Enrichment analyses of 113 tissue-based gene sets across 31 traits

RSS is not restricted to pathways, and can be applied more generally. Here we use it to assess enrichment among tissue-based gene sets that we define based on gene expression data. Specifically we use RNA sequencing data from the Genotype-Tissue Expression project [80] to define sets of the most “relevant” genes in each tissue, based on expression patterns across tissues. The idea is that enrichment of GWAS signals near genes that are most relevant to a particular tissue may suggest an important role for that tissue in the trait.

A challenge here is how to define “relevant” genes. For example, are the highest expressed genes in a tissue the most relevant, even if the genes is ubiquitously expressed [81]? Or is a gene that is moderately expressed in that tissue, but less expressed in all other tissues, more relevant? To address this we considered three complementary methods to define tissue-relevant genes (Methods). The first method (“highly expressed”, HE) uses the highest expressed genes in each tissue. The second method (“selectively expressed”, SE) uses a tissue-selectivity score designed to identify genes that are much more strongly expressed in that tissue than in other tissues (S. Xi, personal communication). The third method (“distinctively expressed”, DE) clusters the tissue samples and identifies genes that are most informative for distinguishing each cluster from others [82]. This last method yields a list of “relevant” genes for each cluster, but most clusters are primarily associated with one tissue, and so we use this to assign genes to tissues. These methods often yield minimally overlapped gene sets for the same tissue (median overlap proportion: 4%; Supplementary Fig. 28)

Despite the small number of tissue-based gene sets relative to the pathway analyses above, this analysis identifies many strong enrichments. The top enriched tissues vary considerably among traits (Table 2), highlighting the benefits of analyzing a wide range of tissues. In addition, traits vary in which strategy for defining gene sets (HE, SE or DE) yields the strongest enrichment results. For example, HE genes in heart show strongest enrichment for heart rate; SE genes in liver show strongest enrichment for LDL. This highlights the benefits of considering multiple annotation strategies, and suggests that, unsurprisingly, there is no single answer to the question of which genes are most “relevant” to a tissue.

**Table 2.** Top enriched tissue-based gene sets in complex traits. Each tissue-based gene set contains 100 transcribed genes used in the Genotype-Tissue Expression project. For each trait we report the most enriched tissue-based gene set (if any) that has a Bayes factor (BF) greater than 1,000 and has more than two member genes with enrichment *P*_1_ > 0.9. All trait-tissue pairs reported above pass the sanity checks (Supplementary Fig. 17). The corresponding background and enrichment parameter estimates are provided in online results (URLs). *P*_1_: the posterior probability that at least one SNP within ±100 kb of the transcribed region of a given gene has non-zero effect on the target trait. HE: highly expressed. SE: selectively expressed. DE: distinctively expressed. *a*: Multiple tissues show partial membership in “Cluster 1”, including ovary, thyroid, spleen, breast and stomach [82]. *b*: These three BFs are smaller than corresponding BFs that SNPs within ±100 kb of transcribed regions of all genes are enriched (Supplementary Fig. 19).

| Phenotype | Tissue<br>(Method) | $\log_{10}$ BF | Select top driving genes<br>(# of genes with enrichment $P_1 > 0.9$) |
| --- | --- | --- | --- |
| Alzheimer's disease | Adrenal gland (SE) | 45.6 | *APOE, APOC1* (2) |
| Neuroticism | Brain (SE) | 26.3 | *LINGO1, KCNC2* (2) |
| Adult height | Nerve tibial (DE) | 25.2<sup>b</sup> | *PTCH1, SFRP4, FLNB* (59) |
| Crohn's disease | Cluster 1<sup>a</sup> (DE) | 15.4 | *SMAD3, ZMIZ1, NUPR1* (6) |
| Inflammatory bowel disease | Cluster 1<sup>a</sup> (DE) | 15.8 | *SMAD3, ZMIZ1, NUPR1* (10) |
| Ulcerative colitis | Heart (HE) | 7.0 | *PLA2G2A, TCAP, ALDOA* (4) |
| Age at natural menopause | Brain (DE) | 1,053.2 | *BRSK1, PPP1R1B, NPTXR* (6) |
| Coronary artery disease | Brain (DE) | 8.5 | *PSRC1, ZEB2, PTPN11* (3) |
| Fasting glucose | Pancreas (SE) | 2,396.8 | *G6PC2, PDX1, SLC30A8* (5) |
| Fasting insulin | Testis (SE) | 866.7 | *ABHD1, PRR30, C2orf16* (3) |
| Heart rate | Heart (HE) | 4.1 | *MYH6, PLN* (5) |
| High-density lipoprotein | Liver (HE) | 20.2 | *APOA1, APOE, MT1G, FTH1* (10) |
| Low-density lipoprotein | Liver (SE) | 33.4 | *ABCG5, LPA, ANGPTL3, HP* (13) |
| Total cholesterol | Liver (DE) | 56.0 | *APOA1, APOE, HP* (9) |
| Triglycerides | Liver (HE) | 93.2 | *APOA1, APOE, FTH1* (7) |
| Serum urate | Kidney (SE) | 210.8<sup>b</sup> | *SLC17A1, SLC22A11, PDZK1* (7) |
| Haemoglobin (HB) | Whole blood (DE) | 2,078.1 | *HIST1H1E, HIST1H1C* (4) |
| Mean cell HB | Whole blood (DE) | 1,363.0 | *NPRL3, FBXO7, UBXN6* (11) |
| Mean cell volume | Whole blood (DE) | 1,019.6<sup>b</sup> | *UBXN6, RBM38, NPRL3* (11) |
| Packed cell volume | Heart (HE) | 945.4 | *RPL19, TCAP* (2) |
| Red blood cell count | Breast (SE) | 141.7 | *OBP2B, STAC2* (2) |

For some traits, the top enriched results (Table 2) recapitulate previously known trait-tissue connections (e.g. lipids and liver, glucose and pancreas), supporting the potential for our approach to identify trait-relevant tissues. Further, many traits show enrichments in multiple tissues. For example, associations in coronary artery disease are strongly enriched in genes related to both *heart* (SE, BF = 6.6 × 10^7^) and *brain* (DE, BF = 3.5 × 10^8^). The multiple-tissue enrichments highlight the potential for our approach to also produce novel biological insights, which we illustrate through an in-depth analysis of late-onset Alzheimer’s disease (LOAD).

Tissue-based analysis of LOAD identified three tissues with very strong evidence for enrichment (BF>10^30^): liver, brain and adrenal gland. Because of the well-known connection between gene *APOE* and LOAD [83], and the fact that *APOE* is highly expressed in these three tissues (Supplementary Fig. 29), we hypothesized that *APOE* and related genes might be driving these results. To assess this we re-analyzed these strongly enriched gene sets after removing the entire apolipoproteins (APO) gene family from them. Of the three tissues, only liver remains (moderately) enriched after excluding APO genes (Fig. 6), suggesting a possible role for non-APO liver-related genes in the etiology of LOAD.

To identify additional genes underlying the liver enrichment, we performed prioritization analysis for non-APO liver-related genes. This highlighted an association of LOAD with gene *TTR* (baseline *P*_1_: 0.64; enrichment *P*_1_: 1.00; Supplementary Fig. 30). *TTR* encodes transthyretin, which has been shown to inhibit LOAD-related protein from forming harmful aggregation and toxicity [84, 85]. Indeed, transthyretin is considered a biomarker for LOAD: patients show reduced transthyretin levels in plasma [86] and cerebrospinal fluid [87]. Rare variants in *TTR* have recently been found to be associated with LOAD [88, 89]. By integrating GWAS with expression data our analysis identifies association of LOAD with *TTR* based on common variants.

## Discussion

We have presented RSS, a new method for simultaneous enrichment and prioritization analysis of GWAS summary data, and illustrated its potential to yield novel insights by extensive analyses involving 31 phenotypes and 4,026 gene sets. We have space to highlight only select findings, and expect that researchers will find the full results (URLs) to contain further insights.

Enrichment tests, sometimes known as “competitive tests” [5, 6], have several advantages over alternative approaches – sometimes known as “self contained tests” (e.g. [90, 91]) – that simply test whether a SNP set contains at least one association. For example, for complex polygenic traits any large pathway will likely contain at least one association, making self-contained tests unappealing. Enrichment tests are also more robust to confounding effects such as population stratification, because confounders that affect the whole genome will generally not create artifactual enrichments. Indeed, in this sense enrichment results can be more robust than single-SNP results. (Nonetheless, most of the summary data analyzed here were corrected for confounding; see Supplementary Table 5.)

**Fig 6:**
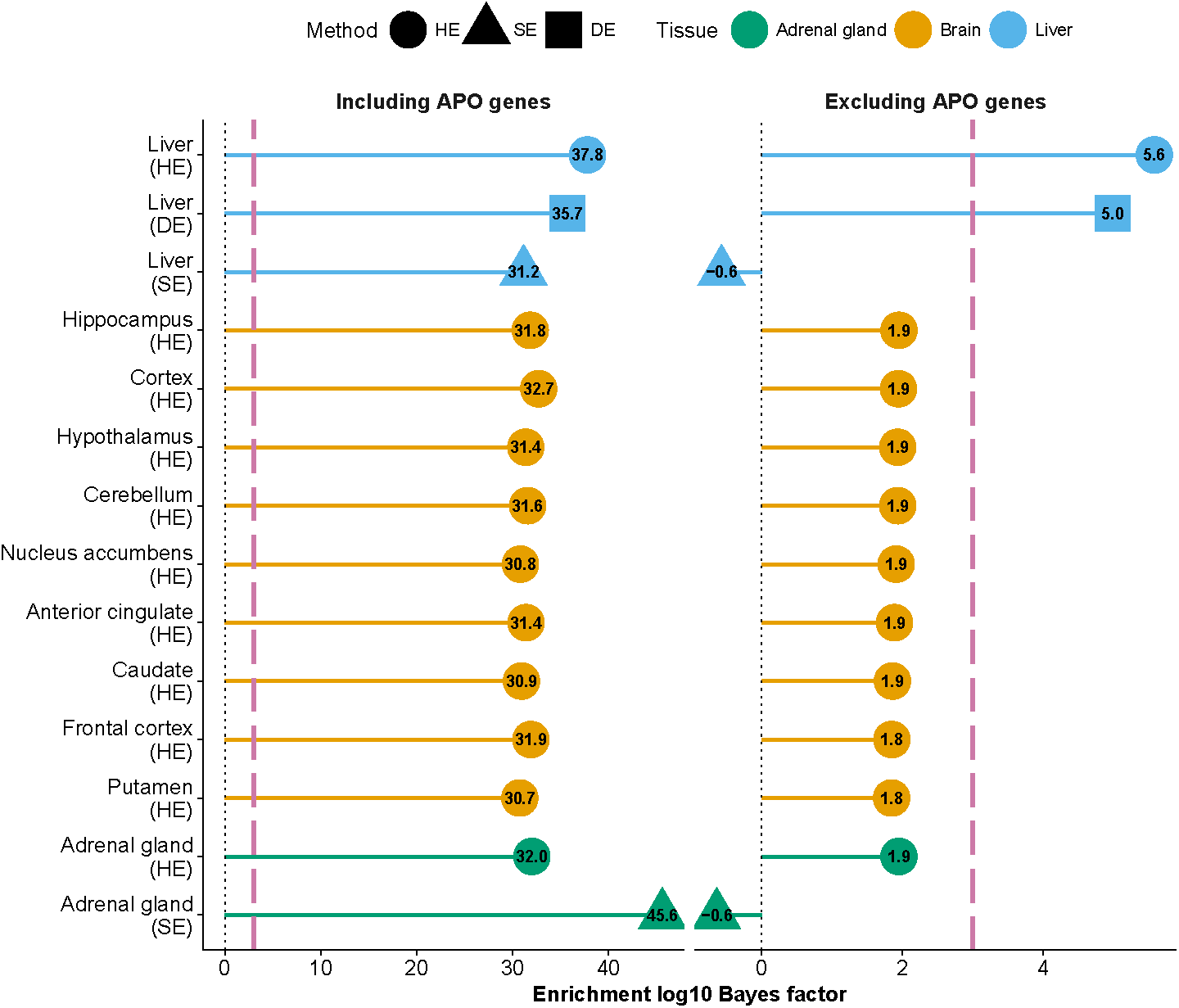
Enrichment analyses of genes related to liver, brain and adrenal gland for Alzheimer’s disease. Shown are the tissue-based gene sets with the strongest enrichment signals for Alzheimer’s disease. Each gene set was analyzed twice: the left panel corresponds to the analysis based on the original gene set; the right panel corresponds to the analysis where SNPs within ± 100 kb of the transcribed region of any gene in Apolipoproteins (APO) family (URLs) are excluded from the original gene set. Dashed lines in both panel denote the same Bayes factor threshold (1,000) used in our tissue-based analysis of all 31 traits (Table 2). HE: highly expressed. SE: selectively expressed. DE: distinctively expressed.

Compared with other enrichment approaches, RSS has several particularly attractive features. First, unlike many methods (e.g. [5, 92, 93]) RSS uses data from *all* variants, and not only those that pass some significance threshold. This increases the potential to identify subtle enrichments even in GWAS with few significant results. Second, RSS models enrichment directly as an increased rate of association of variants within a SNP set. This contrasts with alternative two-stage approaches (e.g. [15, 94, 95]) that first collapse SNP-level association statistics into gene-level statistics, and then assesses enrichment at the gene level. Our direct modeling approach has important advantages, most obviously that it avoids the difficult and error-prone steps of assigning SNP associations to individual genes, and collapsing SNP-level associations into gene-level statistics. For example, simply assigning SNP associations to the nearest gene may highlight the “wrong” gene and miss the “correct” gene [55]. Although our enrichment analyses of gene sets do involve assessing proximity of SNPs to genes in each gene set, they *avoid uniquely assigning each SNP to a single gene*, which is a subtle but important distinction. Finally, and perhaps most importantly, our model-based enrichment approach leads naturally to prioritization analyses that highlight which genes in an enriched pathways are most likely to be trait-associated. We know of only two published methods [8, 96] with similar features, but both require individual-level data and so could not perform the analyses presented here.

Although previous studies have noted potential benefits of integrating gene expression with GWAS data, our enrichment analyses of expression-based gene sets are different from, and complementary to, this previous work. For example, many studies have used expression quantitative trait loci (eQTL) data to help inform GWAS results (e.g. [97–104]). In contrast we bypass the issue of detecting (tissue-specific) eQTLs by focusing only on differences in gene expression levels among tissues. And, unlike methods that attempt to (indirectly) relate expression levels to phenotype (e.g. [105, 106]), our approach focuses firmly on genotype-phenotype associations. Nonetheless, as our results from different tissue-based annotations demonstrate, it can be extremely beneficial to consider multiple approaches, and we view these methods as complimentary rather than competing.

Like any method, RSS also has limitations that need to be considered when interpreting results. For example, annotating variants as being “inside a gene set” based on proximity to a relevant gene, while often effective, can occasionally give misleading results. We saw an example of this when our method identified an enrichment of SE genes in testis with both total cholesterol and triglycerides. Further prioritization analysis revealed that this enrichment was driven by a single gene, *C2orf16* which is a) uniquely expressed in testis, and b) physically close (53 kb) to another gene, *GCKR*, that is strongly associated with lipid traits (Supplementary Fig. 31). This highlights the need for careful examination of results, and also the utility of our prioritization analyses. Generally we view enrichments that are driven by a single gene as less reliable and useful than enrichments driven by multiple genes; indeed, enrichments driven by a single gene seem better represented as a gene association than as a gene set enrichment. Other problems that can affect enrichment methods (not only ours) include: a) an enrichment signal in one pathway can be caused by overlap with another pathway that is genuinely involved in the phenotype; and b) for some traits (e.g. height), genetic associations may be strongly enriched near all genes, which will cause many pathways to appear enriched.

Other limitations of RSS stem from its use of variational inference for approximate Bayesian calculations. Although these methods are computationally convenient in large datasets, and often produce reliable results (e.g. [8, 10, 107–115]), they also have features to be aware of. One feature is that when multiple SNPs in strong LD are associated with a trait, the variational approximation tends to select one of them and ignore the others. This feature will not greatly affect enrichment inference provided that SNPs that are in strong LD tend to have the same annotation (because then it will not matter which SNP is selected). And this holds for the gene-based annotations in the present study. However, it would not hold for “finer-scale” annotations (e.g. appearance in a DNase peak), and so in that setting the use of the variational approximation may need more care. More generally the accuracy of the variational approximation can be difficult to assess, especially since the underlying coordinate ascent algorithm only guarantees convergence to a local optimum. This said, the main alternative for making Bayesian calculations, Markov chain Monte Carlo, can experience similar difficulties.

Finally, the present study examines a single annotation (e.g. one gene set) at a time. Extending RSS to jointly analyze multiple annotations like [14] could further increase power to detect novel associations, and help distinguish between competing correlated annotations (e.g. overlapping pathways) when explaining observed enrichments.

**URLs**. Software, https://github.com/stephenslab/rss; Demonstration of software, http://stephenslab.github.io/rss/Example-5; Full results, https://xiangzhu.github.io/rss-gsea/; APO gene family: http://www.genenames.org/cgi-bin/genefamilies/set/405; Pascal, https://www2.unil.ch/cbg/index.php?title=Pascal;LDSC, https://github.com/bulik/ldsc; COMBAT, https://cran.r-project.org/web/packages/COMBAT; corrplot, https://cran.r-project.org/web/packages/corrplot.

## Acknowledgments

This study is supported by the Grant GBMF # 4559 from the Gordon and Betty Moore Foundation and NIH Grant HG02585 to This study makes use of data generated by the Wellcome Trust Case Control Consortium. A full list of the investigators who contributed to the generation of the data is available from www.wtccc.org.uk. We thank all the GWAS consortia for making their summary statistics publicly available. A full list of acknowledgments is in Supplementary Note. We thank the GTEx consortium for making their RNA sequencing data publicly available. We thank P. Carbonetto and X. He for helpful comments on a draft manuscript. We thank C. A. Anderson, J. R. B. Perry, R. J. F. Loos and M. den Hoed for answering questions related to summary data of specific traits. We thank M. Turchin for downloading summary statistics of red blood cell phenotypes under accession number EGAS00000000132. We thank S. Xi and K. K. Dey for providing expression-based gene lists. This work was completed with resources provided by the University of Chicago Research Computing Center.

## Author Contributions

X.Z. and M.S. conceived the study. X.Z. and M.S. developed the methods. X.Z. developed the algorithms, implemented the software and performed the analyses. X.Z. and M.S. wrote the manuscript.

## Methods

**GWAS summary statistics, LD estimates and SNP annotations**. We analyze GWAS summary statistics of 31 traits, in particular, the estimated single-SNP effect and standard error for each SNP. Following [14], we use the same set of HapMap3 SNPs [16] for all 31 traits, even though some traits have summary statistics available on all 1000 Genomes SNPs [17]. We use this “SNP subsetting” strategy to reduce computation, since the computational complexity of RSS (per iteration) is linear with the total number of SNPs (Supplementary Notes).

Among the HapMap3 SNPs, we also exclude SNPs on sex chromosomes, SNPs with minor allele frequency less than 1%, SNPs in the major histocompatibility complex region, and SNPs measured on custom arrays (e.g. Metabochip, Immunochip) from analyses. The final set of analyzed variants consists of 1.1 million SNPs (Supplementary Table 1, Supplementary Fig. 8).

Since GWAS summary statistics used here were all generated from European ancestry cohorts, we use haplotypes of individuals with European ancestry from the 1000 Genomes Project, Phase 3 [17] to estimate LD [11].

To create SNP-level annotations for a given gene set, we use a distance-based approach from previous enrichment analyses [8, 94]. Specifically, we annotate each SNP as being “inside” a gene set if it is within ± 100 kb of the transcribed region of a gene in the gene set. The relatively broad region is chosen to capture signals from nearby regulatory variants, since the majority of GWAS hits are non-coding.

**Biological pathways and genes**. Biological pathway definitions are retrieved from nine databases (BioCarta, BioCyc, HumanCyc, KEGG, miR-TarBase, PANTHER, PID, Reactome, WikiPathways) that are archived by four repositories: Pathway Commons (version 7) [39], NCBI Biosystems [40], PANTHER (version 3.3) [116] and BioCarta (used in [8]). Gene definitions are based on *Homo sapiens* reference genome GRCh37. Both pathway and gene data were downloaded on August 24, 2015. We use the same protocol described in [8] to compile a list of 3,913 pathways that contains 2-500 autosomal protein-coding genes for the present study. We summarize pathway and gene information in Supplementary Figures 15-16.

**Tissue-based gene sets derived from transcriptome**. Complex traits are often affected by multiple tissues, and it is not obvious *a priori* what the most relevant tissues are for the trait. Hence, it is necessary to examine a comprehensive set of tissues. The breadth of tissues in Genotype-Tissue Expression (GTEx) project [80] provides such an opportunity.

Here we use RNA sequencing data to create 113 tissue-based gene sets. Due to the complex nature of extracting tissue relevance from sequencing data, we consider three different methods to derive tissue-based gene sets.

The first method (“highly expressed”) ranks the mean Reads Per Kilobase per Million mapped reads (RPKM) of all genes based on data of a given tissue, and then selects the top 100 genes with the largest mean RPKM values to represent the target tissue. We downloaded gene lists of 44 tissues with sample sizes greater than 70 from the GTEx Portal on November 21, 2016.

The second method (“selectively expressed”) computes a tissue-selectivity score in a given tissue for each gene, which is essentially the average log ratio of expressions in the target tissue over other tissues, and then uses the top 100 genes with the largest tissue-selectivity scores to represent the target tissue. We obtained gene lists of 49 tissues from S. Xi on February 13, 2017.

The third method (“distinctively expressed”) summarizes 53 tissues as 20 biologically-distinct clusters using admixture models, computes a cluster-distinctiveness score in a given cluster for each gene, and then uses the top 100 genes with the largest cluster-distinctiveness scores to represent the target cluster [82]. We extracted gene lists of 20 clusters from [82] on May 19, 2016.

**Bayesian statistical models**. Consider a GWAS with *n* unrelated individuals typed on *p* SNPs. For each SNP *j*, we denote its estimated single-SNP effect size and standard error as 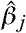 and *ŝ_j_* respectively. To model {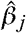, *ŝ_j_*}, we use the **R**egression with **S**ummary **S**tatistics (RSS) likelihood [9]:

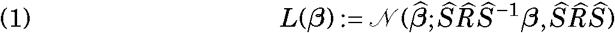
where 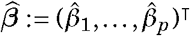, *Ŝ* := diag(**ŝ**), **ŝ**:= (*ŝ*_1_,…,*ŝ_p_*)^⊺^, 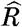 is the LD matrix estimated from an external reference panel with ancestry matching the GWAS cohort, *β* := (*β*_1_,…,*β_p_*)^⊺^ are the true effects of each SNP on phenotype, and 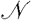 denotes the multivariate normal distribution.

To model enrichment of genetic associations within a given gene set, we borrow the idea from [8] and [56], to specify the following prior on *β*:

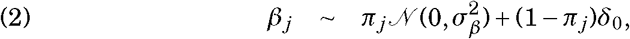

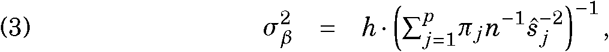

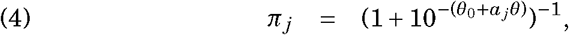
where *δ*_0_ denotes point mass at zero, *θ*_0_ reflects the background proportion of trait-associated SNPs, *θ* reflects the increase in probability, on the log10-odds scale, that a SNP inside the gene set has non-zero effect, *h* approximates the proportion of phenotypic variation explained by genotypes of all available SNPs, and *a_j_* indicates whether SNP *j* is inside the gene set. Following [8], we place independent uniform grid priors on the hyper-parameters {*θ*_0_,*θ*, *h*} (Supplementary Tables 6-7). (If one had specific information about hyper-parameters in a given application then this could be incorporated here.)

**Posterior computation**. We combine the likelihood function and prior distribution above to perform Bayesian inference. The posterior computation procedures largely follow those developed in [10]. First, for each set of hyper-parameters {*θ*_0_,*θ*, *h*} from a predefined grid, we approximate the (conditional) posterior of *β* using a variational Bayes algorithm. Next, we approximate the posterior of {*θ*_0_,*θ*, *h*} by a discrete distribution on the predefined grid, using the variational lower bounds from the first step to compute the discrete probabilities. Finally, we integrate out the conditional posterior of *β* over the posterior of {*θ*_0_,*θ*, *h*} to obtain the full posterior of *β*.

Following [8], we set random initialization as a default for the variational Bayes algorithm. Specifically, we randomly select an initialization, and then use this same initial value for all variational approximations over the grid of {*θ*_0_,*θ*, *h*}. This simple approach was used in all simulations and data analyses for the present study, and yielded satisfying results in most cases.

To facilitate large-scale analyses, we employ several computational tricks. First, we use squared iterative methods [12] to accelerate the fixed point iterations in the variational approximation. Second, we exploit the banded LD matrix [11] to parallelize the algorithm. Third, we use a simplification in [8] that scales the enrichment analysis to thousands of gene sets by reusing expensive genome-wide calculations. See Supplementary Note for details.

For one trait, the total computational cost of our analyses is determined by the number of whole-genome SNPs, the number of gene sets and the grid size for hyper-parameters, all of which can vary considerably among studies. It is thus hard to make general statements about computational time. However, to give a specific example, we finished baseline and enrichment analyses of 1.1 million HapMap3 SNPs and 3,913 pathways for LDL within 36 hours in a standard computer cluster (48 nodes, 12-16 CPUs per node).

All computations in the present study were performed on a Linux system with multiple (4-22) Intel E5-2670 2.6GHz, Intel E5-2680 2.4GHz or AMD Opteron 6386 SE processors.

**Assess gene set enrichment**. To assess whether a gene set is enriched for genetic associations with a target trait, we evaluate a Bayes factor (BF):

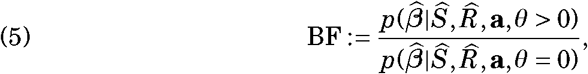
where **a** := (*a*_1_,…, *a_p_*)^⊺^ and *a_j_* indicates whether SNP *j* is inside the gene set. The observed data are BF times more likely under the enrichment model (*θ* > 0) than under the baseline model (*θ* = 0), and so the larger the BF, the stronger evidence for gene set enrichment. See Supplementary Note for details of computing enrichment BF.

**Detect association between a locus and a trait**. To identify trait-associated loci, we consider two statistics derived from the posterior distribution of *β*. The first statistic is *P*_1_, the posterior probability that at least 1 SNP in the locus is associated with the trait:

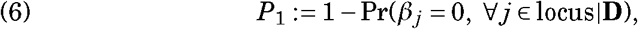
where **D** is a shorthand for the input data including GWAS summary statistics, LD estimates and SNP annotations (if applicable). The second statistic is ENS, the posterior expected number of associated SNPs in the locus:

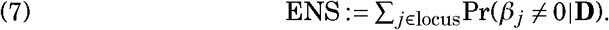

See Supplementary Note for details of computing *P*_1_ and ENS.

**Estimate pairwise sharing of pathway enrichments**. To capture the “sharing” of enrichments between two traits, we define *π* = (*π*_00_,*π*_01_,*π*_10_,*π*_11_):

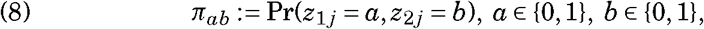
where *z_ij_* equals one if pathway *j* is enriched in trait *i* and zero otherwise. Assuming independence among pathways and phenotypes, we estimate *π* by

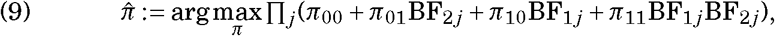
where BF*_ij_* is the enrichment BF for trait *i* and pathway *j*. We solve this optimization problem using an expectation-maximization algorithm implemented in the package ashr [117]. Finally, the conditional probability that a pathway is enriched in a pair of traits given that it is enriched in at least one trait, as plotted in Figure 4b, is estimated as 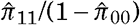.

**Connection with enrichment analysis of individual-level data**. RSS has close connection with the method developed for individual-level data [8]. Under certain conditions [9], we can show that these two methods are mathematically equivalent, in the sense that they have the same fix point iteration scheme and lower bound in variational approximations. See Supplementary Note for proofs. In addition to their theoretical connections, we also empirically compared two methods through simulations, and observed similar inferential results (Supplementary Fig. 32).

**Code and data availability**. Links to source codes and full results of the present study are provided in URLs. Links to GWAS summary statistics are provided in Supplementary Note. HapMap3 SNP list: https://data.broadinstitute.org/alkesgroup/LDSCORE/w_hm3.snplist.bz2. 1000 Genomes Phase 3 data: ftp://ftp.1000genomes.ebi.ac.uk/vol1/ftp/release/20130502. Pathway Commons: http://www.pathwaycommons.org/archives/PC2/v7. NCBI Biosystems: ftp://ftp.ncbi.nih.gov/pub/biosystems. PANTHER: ftp://ftp.pantherdb.org/pathway. BioCarta: https://github.com/pcarbo/bmapathway/tree/master/data. GTEx Portal: https://www.gtexportal.org/home/. “Distinctively expressed” genes [82]: http://stephenslab.github.io/count-clustering.ashr: https://cran.r-project.org/web/packages/ashr.

